# Linking Neural and Clinical Measures of Glaucoma with Diffusion Magnetic Resonance Imaging (dMRI)

**DOI:** 10.1101/370643

**Authors:** Nathaniel Miller, Yao Liu, Roman Krivochenitser, Bas Rokers

## Abstract

**Abstract:** *Purpose:* To link optic nerve (ON) structural integrity to clinical markers of glaucoma using advanced, semi-automated diffusion weighted imaging (DWI) tractography methods in human glaucoma patients.

*Methods:* We characterized optic neuropathy in patients with unilateral advanced-stage glaucoma (n = 6) using probabilistic DWI tractography and compared their results to those in healthy controls (n=6).

*Results:* We successfully identified the ONs of glaucoma patients based on DWI in all patients and confirmed that the degree of reduced structural integrity of the ONs determined using DWI correlated with clinical markers of glaucoma severity. Specifically, we found reduced fractional anisotropy (FA), a measure of structural integrity, in the ONs of eyes with advanced, as compared to mild, glaucoma (F(1,10) = 55.474, p < 0.0001). Furthermore, by comparing the ratios of ON FA in glaucoma patients to those of healthy controls (n = 6), we determined that this difference was beyond that expected from normal anatomical variation (F(1,9) = 20.276, p < 0. 005). Finally, we linked the DWI-measures of neural integrity to standard clinical glaucoma measures. ON vertical cup-to-disc ratio (vCD) predicted ON FA (F(1,10) = 11.061, p < 0.01, R^2^ = 0.66), retinal nerve fiber layer thickness (RNFL) predicted ON FA (F(1,10) = 11.477, p < 0.01, R^2^ = 0.63) and ON FA predicted perceptual deficits (visual field index [VFI]) (F(1,10) = 15.308, p < 0.005, R^2^ = 0.52).

*Conclusion:* We provide semi-automated methods to detect glaucoma-related structural changes using DWI and confirm that they correlate with clinical measures of glaucoma.

## Introduction

Vision loss is a major cause of disability worldwide that is particularly common among the elderly, conferring a greater risk of injury and diminished quality of life [1-2]. Glaucoma is the leading cause of irreversible vision loss worldwide and is projected to affect nearly 80 million individuals by 2020 [3]. It is clinically defined by characteristic patterns of visual field impairment and optic nerve (ON) damage [4]. There is growing evidence that glaucoma may be a neurodegenerative condition. Recent studies suggest that the pathogenesis of glaucoma bears similarities to that of Alzheimer’s disease, Parkinson’s, and amyotrophic lateral sclerosis (ALS) [6]. Thus, research seeking to understand the possible underlying neurodegenerative processes associated with glaucoma may help guide the development of more robust treatment paradigms and have applications for improving our understanding of other neurodegenerative diseases. In addition, current glaucoma treatments are limited to the reduction of intraocular pressure (IOP) to prevent progressive visual field loss. However, many patients continue to lose vision despite treatment, and no treatments are available to reverse damage that has already occurred to the visual system [4]. Therefore, the development of robust methods enabling earlier diagnosis and more precise quantification of disease progression are essential to limiting glaucomatous damage and improving patient outcomes.

MRI-based *in vivo* human studies using voxel-based morphometry (VBM) and diffusion tensor imaging (DTI) to explore gray- and white-matter cortical changes associated with glaucoma have shown reduced gray-matter (GM) volume in late stages of the disease, along with significant rarefaction along the optic radiations [7-9]. While these results are consistent with animal models and post-mortem pathological studies of human subjects [10], there is a need for more precise evaluation of neurological changes, particularly at the level of the ON. Animal model studies of the early visual system using diffusion MRI (dMRI) demonstrate the ability to detect changes in the neural integrity of the ONs from damage occurring within the retina [11-12]. In a meta-analysis of studies examining the ONs of human glaucoma patients using various DTI methods, Li, et al. (2014) noted significant decreases in fractional anisotropy (FA) and increases in mean diffusivity (MD) in the ONs of glaucoma patients compared to controls [13]. A number of studies in this meta-analysis also examined the correlation between various measures of disease severity (including glaucoma stage and optical coherence tomography [OCT] measurements) and structural white-matter changes. Generally, increasing glaucoma disease severity is associated with greater white-matter disruption (i.e. decreasing FA and increasing MD) [14-18].

While these studies have quantified the diffusion properties of glaucomatous ONs and their relationship with various clinical measures of disease severity, they rely on older dMRI methodologies. In particular, these studies employ relatively low-resolution, single phase-encoding direction diffusion scans and sample the ONs using manually placed regions of interest (ROIs). [14-19]. By sampling from only small portions of the ONs, these techniques are limited in their ability to measure the full extent of optic neuropathy. Moreover, manual ON segmentation is time-intensive, may introduce operator error, and limits the ability to translate this technique into widespread clinical practice. Diffusion MRI methods have advanced substantially since the publication of these studies, and there is an opportunity to investigate and validate methods that rely on semi-automated techniques to assess the ON using dMRI.

Recent diffusion-weighted imaging (DWI) based probabilistic tractography methods have been developed that can more precisely evaluate white-matter changes in the human visual system. These methods have demonstrated reduced white matter integrity in patients with amblyopia [20]. In this study, we apply this technique to patients with asymmetric glaucomatous optic neuropathy in each eye to evaluate DWI methods as a diagnostic tool for visual disorders, linking changes in white-matter integrity to retinal and perceptual changes in glaucoma. Recent methodological advances make it possible to reliably identify the microstructural properties of the ON with limited user input [21]. Using a pair of diffusion scans acquired with opposite phase-encoding directions, a low-noise field-corrected volume can be created [22-23], allowing the ONs to be isolated using probabilistic tractography. This provides a unique opportunity to quantify changes across the entire length of the ONs in glaucoma patients. Further, we purposefully selected patients with asymmetric glaucomatous ON damage to allow for within-subjects comparisons of ON properties, quantifying differences in eyes with “advanced” versus “mild” glaucoma as defined by the American Academy of Ophthalmology [24].

We used an advanced DWI tractography method to identify and analyze the ONs of six asymmetric glaucoma patients and six controls. Using both within-subject analyses and comparison to controls, we evaluated structural changes in the ONs associated with glaucoma. Furthermore, we assessed the relationship between these MRI-based neural measures, clinical measures of ON and retinal integrity (e.g. vertical cup-to-disc ratio and average peripapillary retinal nerve fiber layer thickness), and perceptual measures (e.g. visual field index).

## Methods

### Participants

Our study was conducted in accordance with the Code of Ethics of the World Medical Association (Declaration of Helsinki). Informed consent was obtained from all participants and all participants completed MRI screening with consultation and approval obtained from their physicians as needed to ensure they could safely participate.

#### Glaucoma Patients

Six glaucoma patients (4 female) aged 19-66 years (mean 53.3 ± 17.4) were recruited from the Glaucoma Service of the University of Wisconsin Hospitals and Clinics (Table 1). All patients had a diagnosis of either primary open-angle, pigment dispersion, pseudoexfoliation, or chronic angle-closure glaucoma and a history of IOPs greater than 22 mmHg. Selection criteria included a Snellen best-corrected visual acuity of 20/25 or better in the eye with “mild” glaucoma and 20/200 or better in the eye with “advanced” glaucoma. Patients with any history of neurodegenerative diseases, normal-tension glaucoma, diabetic retinopathy, advanced macular degeneration, uveitis, or previous (non-surgical) eye trauma were excluded.

**Table 1.** Demographics and Ocular Characteristics of Glaucoma Patients

| Patient | Age | Sex | Eye | VA | VFI | vCD | RNFL (μm) |
| --- | --- | --- | --- | --- | --- | --- | --- |
| G1 | 19 | F | OD | 20/20 | 100% | 0.34 | 121 |
|  |  |  | OS | **20/30** | **41%** | **0.83** | **56** |
| G2 | 54 | M | **OD** | **20/20** | **92%** | **0.66** | **66** |
|  |  |  | OS | 20/20 | 99% | 0.63 | 70 |
| G3 | 57 | F | **OD** | **20/25** | **54%** | **0.82** | **52** |
|  |  |  | OS | 20/30 | 99% | 0.63 | 73 |
| G4 | 59 | M | **OD** | **20/30** | **62%** | **0.82** | **45** |
|  |  |  | OS | 20/20 | 97% | 0.80 | 57 |
| G5 | 65 | F | OD | 20/40 | 96% | 0.57 | 73 |
|  |  |  | OS | **20/25** | **69%** | **0.78** | **52** |
| G6 | 66 | F | OD | 20/20 | 100% | 0.74 | 83 |
|  |  |  | OS | **20/20** | **91%** | **0.89** | **66** |

Characteristics of glaucoma patients including age, sex, Snellen best-corrected visual acuity (VA), visual field index (VFI), vertical cup-to-disc ratio (vCD), and average peripapillary retinal nerve fiber layer thickness (RNFL). Eyes with advanced glaucoma are indicated in **bold**.

#### Control Subjects

Control subjects were recruited from the University of Wisconsin-Madison. Six gender-matched subjects aged 21-34 years (mean 24 ± 5.3) were included in the analysis. All subjects had Snellen best-corrected visual acuity of 20/20 or better and had no prior medical history of neurologic or ocular pathology other than refractive error. Eye dominance was determined as follows: subjects were instructed to form a small aperture using both hands (right and left hands overlapping so a small opening is formed with the inner sides of the palms and thumbs) and to fixate on a distant object through that opening with both eyes open. Without moving their head or hands, subjects were then instructed to close their left eye and were asked whether or not they could still see the object. This same task was repeated with the right eye closed. The eye with which they could see the fixation target was recorded as the “dominant” eye. Ocular dominance was successfully determined for 6/6 control subjects.

### Clinical Measures

Clinical measures of ON structure and function were assessed for each of the six glaucoma patients during clinical ophthalmologic exams by a glaucoma specialist (Y.L.). These measures included Snellen best-corrected visual acuity (VA), vertical cup-to-disc ratio (vCD), average peripapillary retinal nerve fiber layer thickness (RNFL), and visual field index (VFI). The vCD was determined by direct visualization of the ON using slit-lamp biomicroscopy, average peripapillary RNFL thickness was measured using Cirrus Spectral-Domain Optical Coherence Tomography (Carl Zeiss Meditec, Inc., Dublin, CA, USA) with all scans having adequate signal strength (>7/10), and VFI was measured using the Humphrey visual field 24-2 SITA-Standard testing algorithm (Carl Zeiss Meditec, Inc. Dublin, CA, USA) on visual fields with adequate reliability indices (<33% fixation losses, false positives, and false negatives).

### Magnetic Resonance Imaging Data Acquisition

Brain imaging data was obtained at the Waisman Center in Madison, WI using a GE Discovery Medical 3T MRI scanner (GE Healthcare, Inc., Chicago, IL, USA) equipped with a 32-channel head coil. First, a 10-minute structural whole-brain T1-weighted anatomical scan (2.93 ms TE; 6.70 ms TR; 1 mm^3^ isotropic voxels) was acquired. Then, a 15-minute diffusion sequence with two 48-direction diffusion-weighted scans (6 b0), collected in the anterior to posterior (AP) and posterior to anterior (PA) directions (76.7 ms TE; 8.1 s TR; 2×2×2 mm^3^ isotropic voxels; b = 2000 nm/s^2^; reconstruction matrix FOV: LR 212 mm x AP 212 mm x FH 144 mm).

### Data Processing

#### Pre-processing

To improve the quality of our tractography and increase the signal-to-noise ratio in the nasal cavity, the two reverse-encoded (AP and PA) diffusion scans were combined into a single corrected volume using the FSL software (University of Oxford, Oxford, England) [22-23].

Subsequent processing was completed using the mrVista software package (Stanford University, Stanford, California), based on previously published methods [21]. A mean *b=0* image was calculated from the corrected DTI volume and underwent eddy current correction. This corrected b0 image was co-registered to the AC-PC-aligned T1 image and diffusion tensors were fit to the volume using a least-squares estimate bootstrapped 500 times [25].

#### ROI Placement

We manually identified three ROIs along the brain’s visual pathway. The T1 image was used to place the left and right ONs and the optic chiasm (OC) by gross anatomy. 4-mm spheres were used for the ONs (centered slightly posterior to the ON head at the back of the eye), and a 6-mm sphere was used for the OC.

#### Tractography

We derived visual pathways through probabilistic diffusion-weighted tractography using MRtrix2 (Brain Research Institute, Melbourne, Australia) [26-34]. Constrained spherical deconvolution (CSD) estimates were used to generate fibers between two ROI pairs, representing the left and right ONs (ON » OC). Whole-brain tractography was completed using an *L_max_* of 6 with 500,000 seeds and a maximum of 5,000,000 fibers. A modified white-matter mask generated using mrVista was used to constrain fibers to the brain while still allowing CSDs to be fit within the nasal cavity, enabling detection of the ONs. Final pathways were restricted to fibers passing between the specified ROIs, omitting any spurious results.

#### Fiber Cleaning

Fiber groups were cleaned using the Automated Fiber Quantification (AFQ) toolkit (Stanford University) [35], followed by a quality assessment and manual cleaning as necessary. Fibers were overlaid on the anatomical T1 volume and any fibers that were found to be anatomically implausible were manually removed. Pathways from all 12 participants were processed using the same AFQ cleaning parameters and were manually refined by the same operator (N.M.) (Fig. 1).

**Fig 1.**
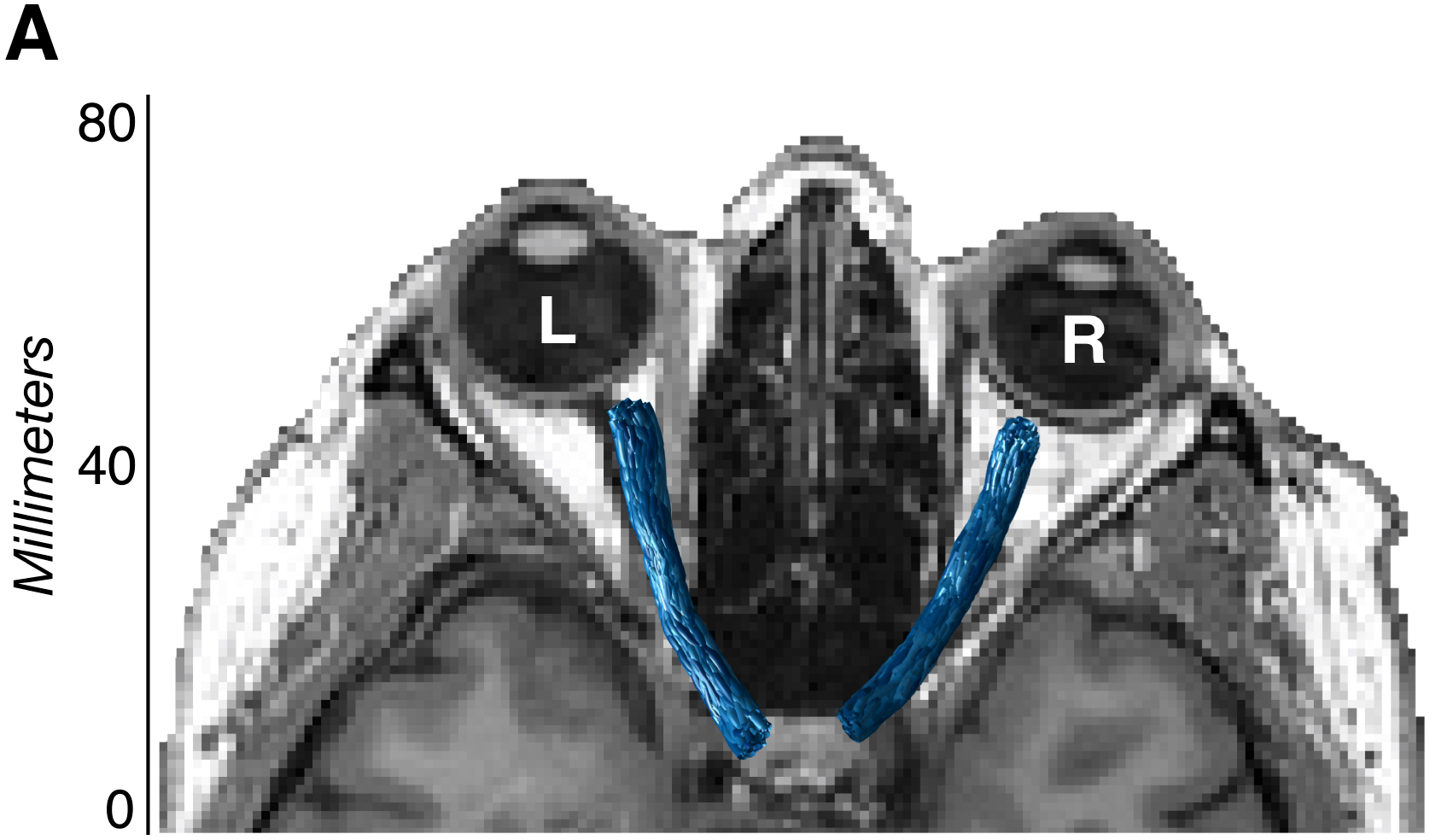
Optic nerve visualization using diffusion-weighted magnetic resonance imaging. Visualization of final tractography-generated optic nerve white-matter pathways (blue) in a representative glaucoma patient (G6) using diffusion-weighted magnetic resonance imaging.

#### Diffusion Measures

Voxel-wise tensor properties were extracted from the volumetric region defined by each tractography-generated pathway. The main diffusion properties included in our analysis were mean diffusivity (MD, μm^2^/s) and fractional anisotropy (FA) (Fig 2). MD provides an average measure of pathway diffusivity and is a useful approximation of white-matter density, where large values indicate a diffuse (“weak”) pathway, and small values indicate a denser and/or more myelinated (“strong”) pathway [36]. FA provides a measure of diffusion directionality and is highly sensitive to microstructural changes across different pathologies, where large values indicate a single highly myelinated “intact” pathway, and small values indicate multiple intersecting, degenerated, or demyelinated pathways. To more precisely characterize the MD and FA measures, we also assessed the component measures, radial diffusivity (RD, μm^2^/s) and axial diffusivity (AD, μm^2^/s). RD has been demonstrated to be more sensitive to changes in white-matter myelination, while AD is more sensitive to axonal degeneration [37]. Typically, large RD values indicate demyelination, while large AD values indicate axonal degeneration.

**Fig 2.**
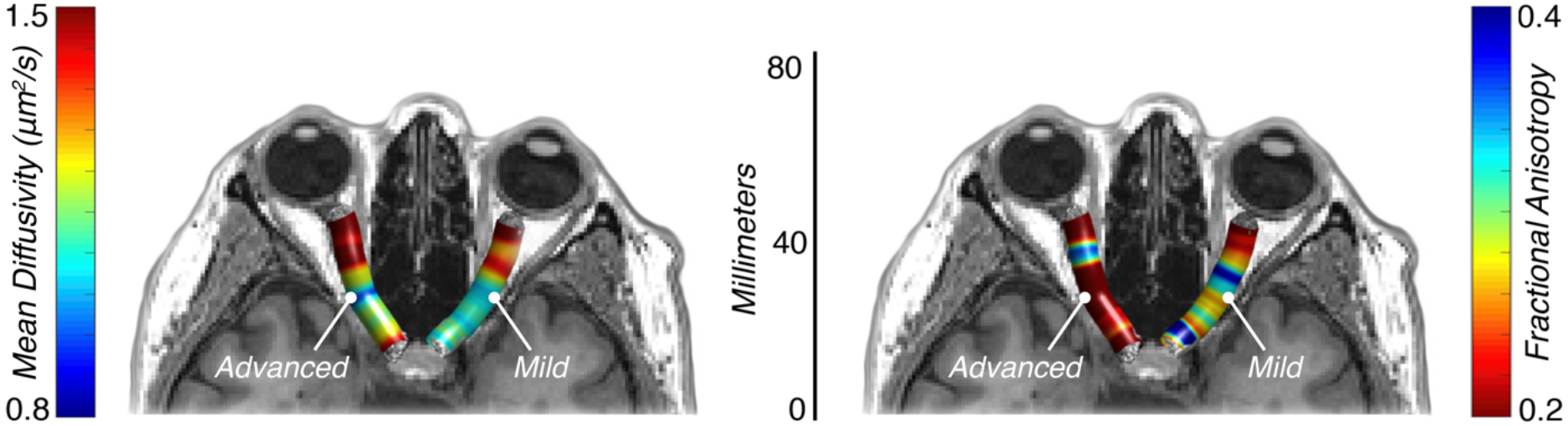
Optic nerve diffusion properties using diffusion weighted magnetic resonance imaging. Mean diffusivity (μm^2^/s) and fractional anisotropy values in a representative glaucoma patient (G6) with advanced glaucoma in the left optic nerve and mild glaucoma in the right optic nerve using diffusion weighted magnetic resonance imaging.

#### Analysis

A combination of within- and between-groups analyses were conducted to evaluate the structural white-matter changes associated with glaucoma-related vision damage. All six asymmetric glaucoma patients had one eye with no or early glaucomatous visual field defects (“mild”) and one eye with moderate or advanced glaucomatous visual field defects (“advanced”) [24]. Within-subjects, diffusion-tensor properties were compared between the “advanced” and “mild” eyes. Between-subjects, the ratios of “advanced” / “mild” ON FA in glaucoma patients were compared to non-dominant/dominant ratios in controls. This design minimized the possible effect of global changes in white-matter properties as a result of age.

To facilitate these comparisons and normalize pathway lengths, 100 samples were taken along the length of each pathway such that 100 average MD, FA, RD, and AD values were available for each pathway in each of the two groups (glaucoma patients and control subjects). These values were generated for each cross-section using a Gaussian-weighted average, where the calculated “core” of each pathway was selectively weighted over the outlying fibers. From these 100 samples, the central 80 were retained for further analysis to reduce the risk of including measures contaminated by retinal cell bodies or contralateral fiber tracts [20-21, 38]. The central 80% of each sample was further subdivided into 10% bins to more precisely quantify differences along the pathway length.

#### Statistical Analysis

A linear mixed effect (LME) model was used to compare the MD, FA, RD, and AD values of the advanced and mild ONs across the middle 80% of samples and at each of the eight 10% bins in glaucoma patients. This model factored glaucoma severity (advanced or mild) as a fixed effect and subject as a random effect. Reported p-values are from ANOVAs of the fixed “glaucoma severity” effects. The same LME model was used in the glaucoma and control ratio comparisons, as well as in all correlational data (factoring clinical and neurological measures as fixed effects and subjects as random effects). For all correlations, reported R^2^ values are adjusted to the number of predictors included in the LME model.

## Results

### Selection of Clinical Measures of Structure and Function

As expected, we identified strong correlations between vCD and RNFL (F(1,10) = 229.17, p = 3.20e-8, R^2^ = 0.98), vCD and VFI (F(1,10) = 15.662, p = 0.0027, R^2^ = 0.64), and RNFL and VFI (F(1,10) = 24.228, p = 0.00060, R^2^ = 0.76) (Fig 3). These measures quantify the anticipated correlations between clinical measures of structural and functional glaucomatous optic nerve damage in our patient sample.

**Fig 3.**
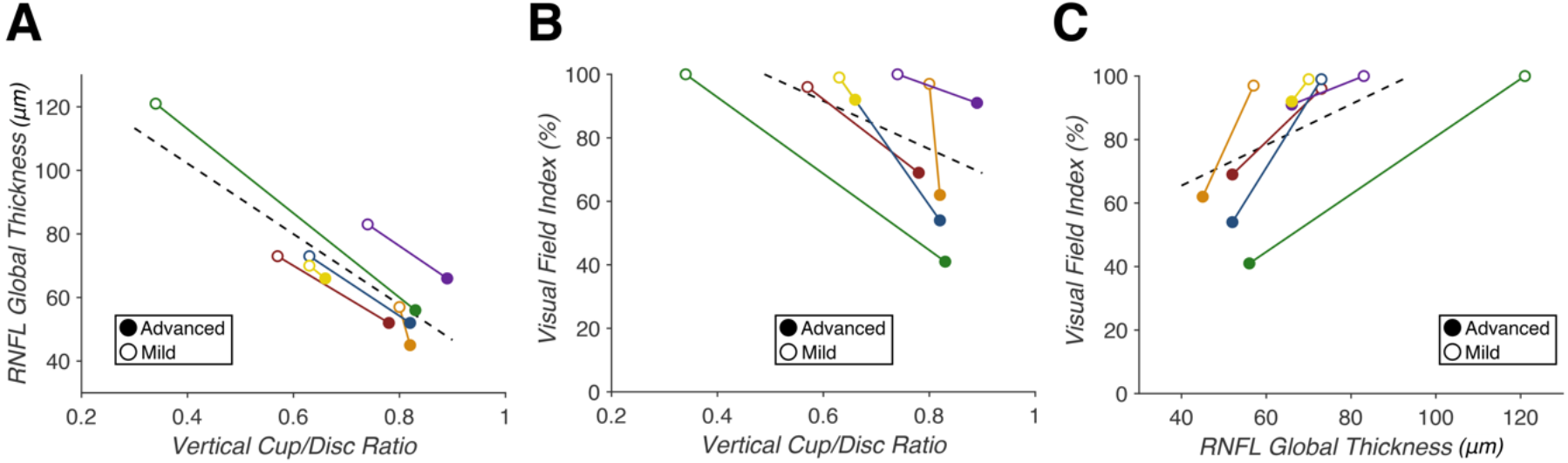
Correlations between clinical glaucoma measures in glaucoma patients with asymmetric optic nerve damage (n=6) (A) Vertical cup/disc ratio (vCD) predicts average retinal nerve fiber layer (RNFL) thickness (p = 3.20e-8, R^2^ = 0.98). (B) vCD predicts visual field index (VFI) (p = 0.0027, R^2^ = 0.64). (C) RNFL predicts VFI (p = 0.00060, R^2^ = 0.76). Correlations within individual patients are indicated by each solid colored line, with closed points marking eyes with “advanced” glaucoma and open points marking eyes with “mild” glaucoma. A least-squares regression estimate is indicated by the dashed line.

### Diffusion Magnetic Resonance Imaging

ON white-matter pathways were successfully identified and refined in 6/6 glaucoma patients and 6/6 controls. All pathways appeared to be anatomically plausible after cleaning and were amenable to within-subjects and group-wise comparisons. A significant difference in mean FA between the advanced and mild ONs of glaucoma patients was noted in 3/8 bins (F_1_(1,10) = 55.442, p_1_ = 2.20e-5; F_2_(1,10) = 18.382, p_2_ = 0.0016; F_3_(1,10) = 11.322, p_3_ = 0.0072), along with a significant difference across the middle 80% of samples (F(1,10) = 55.474, p = 2.19e-5) (Fig 4). In the same pathway, a significant difference in average MD was noted in 1/8 bins (F(1,10) = 10.885, p = 0.0080). For RD, we found a significant difference in 3/8 bins (F_1_(1,10) = 15.87, p_1_ = 0.0026; F_2_(1,10) = 4.974, p_2_ = 0.0498; F_3_(1,10) = 5.687, p_3_ = 0.038), along with a significant difference across the middle 80% (F(1,10) = 5.118, p = 0.047). No significant difference in AD was found (all p > 0.05). Our subsequent analyses focus on ON FA because of the magnitude and reliability of the effect (compared to MD and RD).

**Fig 4.**
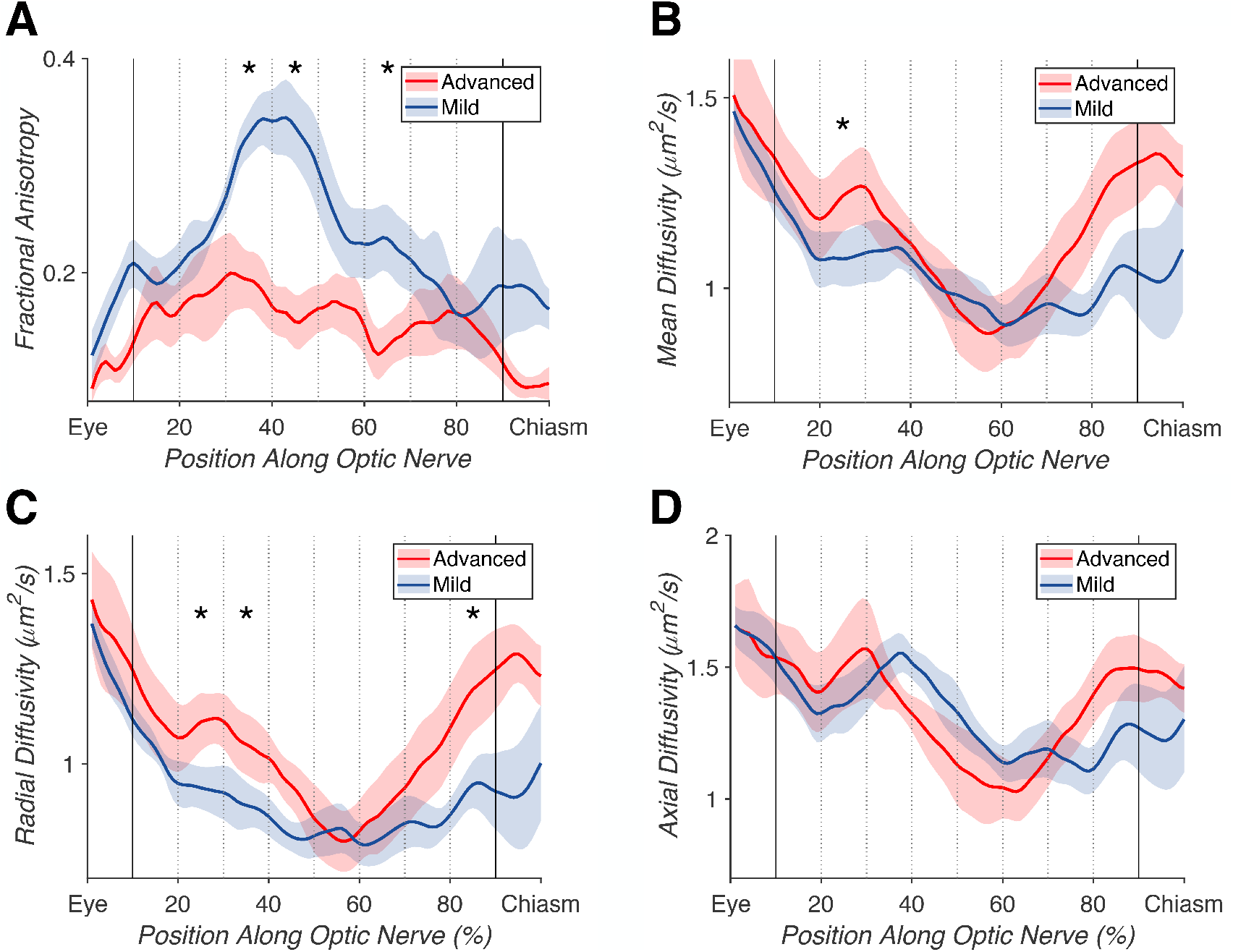
Advanced versus mild glaucomatous optic nerve (ON) tract profiles. Average tract profiles for the ONs of 6 eyes with advanced (red) and 6 eyes with mild (blue) glaucoma, with the middle 80% of samples marked with bold lines, and each 10% bin marked with dotted lines. Significant differences (p < 0.05) denoted by *. (A) Differences in fractional anisotropy across the middle 80% of samples (p = 2.19e-5), and in 3/8 individual bins (p_1_ = 2.20e-5; p_2_ = 0.0016; p_3_ = 0.0072). (B) Difference in mean diffusivity in 1/8 bins (p = 0.0080). (C) Differences in radial diffusivity across the middle 80% (p = 0.047), and in 3/8 individual bins (p_1_ = 0.0026, p_2_ = 0.0498, p_3_ = 0.038). (D) Difference in axial diffusivity was not significant (all p > 0.05).

To more precisely characterize the nature of these within-subject effects, we compared the FA ratios of advanced/mild ONs in glaucoma patients to non-dominant/dominant ONs in controls. We found selective FA reductions in the “advanced” ONs of glaucoma patients. As expected, there were no significant differences between non-dominant and dominant ON FA values in control subjects. We found a significant difference between the ON FA ratios of glaucoma patients compared to controls in a LME model including group and age as fixed factors (F(1,9) = 20.276, p = 0.0015) (Fig 5).

**Fig 5.**
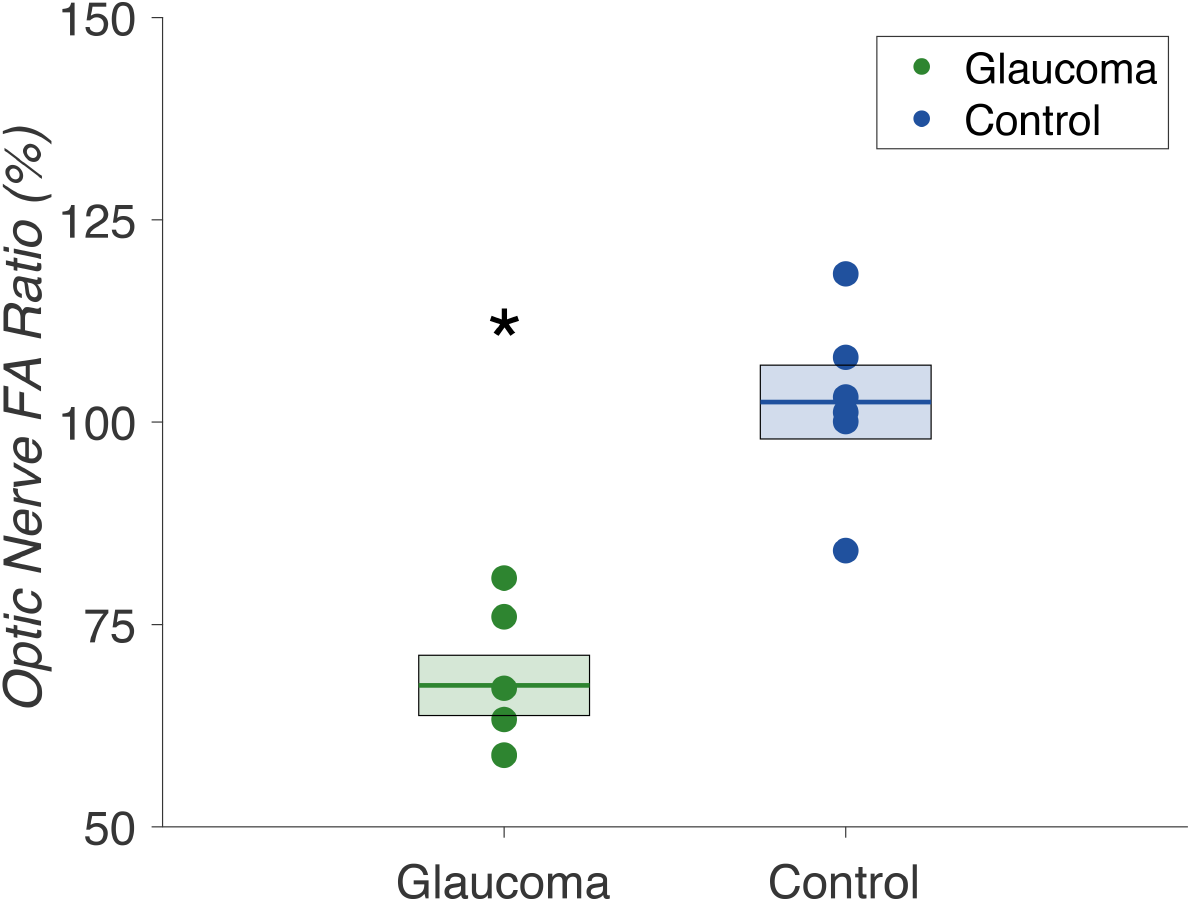
Optic nerve fractional anisotropy ratios in glaucoma patients (n=6) and controls (n=6) Comparison of optic nerve fractional anisotropy ratios (%) in glaucoma patients (green) and controls (blue). Glaucoma patient ratios were calculated for “advanced” / “mild” optic nerve fractional anisotropy, and control subject ratios were calculated for non-dominant/dominant optic nerve fractional anisotropy. Mean ratios are indicated by the bold lines, with standard error denoted by the surrounding box. We found a significant difference between fractional anisotropy ratios in glaucoma patients versus controls (p = 0.0015).

### Relating dMRI to Clinical Measures in Glaucoma

Reductions in ON FA were found to correlate with clinical measures of glaucoma (vCD, RNFL, and VFI). We found that vCD predicted ON FA (F(1,10) = 11.061, p = 0.0077, R^2^ = 0.66), RNFL predicted ON FA (F(1,10) = 11.477, p = 0.0069, R^2^ = 0.63), and ON FA in turn predicted VFI (F(1,10) = 15.308, p = 0.0029, R^2^ = 0.52) (Fig 6). Thus, dMRI measures of white matter integrity in the optic nerve reliably linked the retinal and perceptual deficits observed in our patient sample.

**Fig 6.**
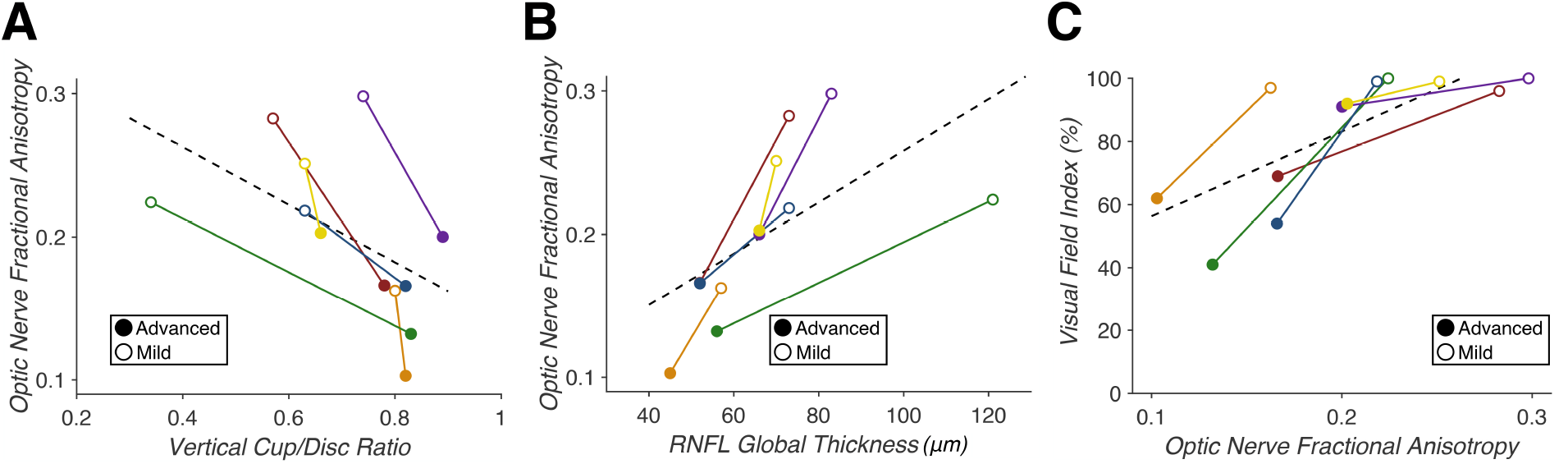
Correlation of dMRI fractional anisotropy with clinical measures of glaucoma (n=6) (A) Vertical cup-to-disc ratio predicts optic nerve fractional anisotropy (p = 0.0077, R^2^ = 0.66). (B) Average retinal nerve fiber layer thickness predicts optic nerve fractional anisotropy (p = 0.0069, R^2^ = 0.63). (C) Optic nerve fractional anisotropy predicts visual field index (p = 0.0029, R^2^ = 0.52). Correlations within individual patients are indicated by each solid colored line, with closed points marking eyes with “advanced” glaucoma and open points marking eyes with “mild” glaucoma. A least-squares regression estimate is indicated by the dashed line.

## Discussion

We assessed the utility of probabilistic DWI tractography in correlating neural and clinical measures of ON damage in patients with asymmetric glaucoma and normal controls. We isolated the ON white-matter pathway in 6/6 glaucoma patients and 6/6 controls. We combined AP and PA phase-encoded dMRI volumes to recover from imaging distortions caused by the adjacent nasal cavities. To our knowledge, this is the first probabilistic tractography study to successfully isolate the ONs in visually-impaired patients. Previous work relied largely on manual ON segmentation or ROI-based analyses, while our methodology allows the entire ON to be identified with minimal manual operator input. This technique increases the efficiency of data collection and reduces the risk of operator error. We correlated clinical measures of ON structure and function (i.e. vCD, RNFL, and VFI) with dMRI measures of ON integrity (i.e. FA, MD, RD, and AD). Our methods sample diffusion measures along the entire length of the ON (rather than small targeted regions) and provides a more comprehensive account of dMRI measures of disease.

We found significant differences in average FA, MD, and RD of ONs with “advanced” versus “mild” glaucoma. These trends (smaller FA and greater MD and RD in “advanced” ONs) indicate reduced integrity of the visual system consistent with clinical measures of glaucoma severity. Our findings are mostly consistent with earlier studies, which showed general trends of decreasing FA and increasing MD with increasing glaucoma severity [14-19]. However, there was a significant difference in radial, but not axial diffusivity, which would suggest that glaucoma predominantly impacted myelination rather than axonal degeneration of ONs with advanced glaucoma. This result differs from previous work, where changes were noted in both RD and AD [16]. This discrepancy may be the result of differences in diffusion sequence parameters but is most likely the result of our small sample size.

To minimize the impact of the small sample size and the difference in mean age between our glaucoma patients and controls in this study, we primarily relied on within-subjects comparisons. Through ratio comparison of ON FA ratios in glaucoma patients to controls, we determined that these structural changes were not global changes in the ONs of glaucoma patients, but rather unilateral changes in the “advanced” glaucomatous eyes.

Lastly, we examined the correlation between clinical glaucoma measures (vCD, RNFL, and VFI) and dMRI neural measures (ON FA). We found that vCD and RNFL were correlated with ON FA and that ON FA was predictive of VFI. Thus, our analysis confirms and quantifies correlations between measures of glaucoma-related damage between neural (ON FA), structural (vCD), retinal (RNFL), and functional (VFI) measures.

In summary, we correlated neural DWI measures and clinical glaucoma measurements in patients with asymmetric glaucoma damage. Our results using current dMRI methods agree with previous studies using older techniques and validate probabilistic tractography methods that assess deficits in the visual pathways of glaucoma patients. Future larger, prospective studies may evaluate dMRI as a possible diagnostic tool for glaucoma evaluation. In addition, studies quantifying the relationship between neural and clinical measures longitudinally may determine whether these methods may be useful for monitoring glaucoma progression and treatment efficacy.

## Acknowledgments

We thank Anna Bauman for assistance with patient recruitment and data collection, as well as Franco Pestilli for comments on a prior version of the manuscript.

## References

1. National Academies of Sciences E, and Medicine. Making Eye Health a Population Health Imperative: Vision for Tomorrow. Teutsch SM, McCoy MA, Woodbury RB, Welp A, editors. Washington, DC: The National Academies Press; 2016. 586 p.

2. Vu HTV, Keeffe JE, McCarty CA, Taylor HR. Impact of unilateral and bilateral vision loss on quality of life. The British Journal of Ophthalmology. 2005;89(3):360–3. doi: 10.1136/bjo.2004.047498. PubMed PMID: PMC1772562.

3. Quigley HA, Broman AT. The number of people with glaucoma worldwide in 2010 and 2020. British Journal of Ophthalmology. 2006;90(3):262–7. doi: 10.1136/bjo.2005.081224. PubMed PMID: WOS:000235424800008.

4. Davis BM, Crawley L, Pahlitzsch M, Javaid F, Cordeiro MF. Glaucoma: the retina and beyond. Acta Neuropathologica. 2016;132(6):807–26. doi: 10.1007/s00401-016-1609-2. PubMed PMID: Davis2016.

5. Leske MC. Open-angle glaucoma - An epidemiologic overview. Ophthalmic Epidemiology. 2007;14(4):166–72. doi: 10.1080/09286580701501931. PubMed PMID: WOS:000249724700004.

6. Jindal V. Glaucoma: An Extension of Various Chronic Neurodegenerative Disorders. Molecular Neurobiology. 2013;48(1):186–9. doi: 10.1007/s12035-013-8416-8. PubMed PMID: Jindal2013.

7. Engelhorn T, Michelson G, Waerntges S, Struffert T, Haider S, Doerfler A. Diffusion Tensor Imaging Detects Rarefaction of Optic Radiation in Glaucoma Patients. Academic Radiology. 2011;18(6):764–9. doi: 10.1016/j.acra.2011.01.014. PubMed PMID: WOS:000290976900015.

8. Frezzotti P, Giorgio A, Toto F, De Leucio A, De Stefano N. Early changes of brain connectivity in primary open angle glaucoma. Human Brain Mapping. 2016;37(12):4581–96.

9. Zikou AK, Kitsos G, Tzarouchi LC, Astrakas L, Alexiou GA, Argyropoulou MI. Voxel-Based Morphometry and Diffusion Tensor Imaging of the Optic Pathway in Primary Open-Angle Glaucoma: A Preliminary Study. American Journal of Neuroradiology. 2012;33(1):128–34. doi: 10.3174/ajnr.A2714. PubMed PMID: WOS:000299491400022.

10. Gupta N, Yucel YH. Glaucoma as a neurodegenerative disease. Curr Opin Ophthalmol. 2007;18(2):110–4. Epub 2007/02/16. doi: 10.1097/ICU.0b013e3280895aea. PubMed PMID: 17301611.

11. Song S-K, Sun S-W, Ju W-K, Lin S-J, Cross AH, Neufeld AH. Diffusion tensor imaging detects and differentiates axon and myelin degeneration in mouse optic nerve after retinal ischemia. NeuroImage. 2003;20(3):1714–22. doi: 10.1016/j.neuroimage.2003.07.005. PubMed PMID: 14642481.

12. Xu J, Sun S-W, Naismith RT, Snyder AZ, Cross AH, Song S-K. Assessing Optic Nerve Pathology with Diffusion MRI: from Mouse to Human. NMR in biomedicine. 2008;21(9):928–40. doi: 10.1002/nbm.1307. PubMed PMID: PMC2603138.

13. Li K, Lu C, Huang Y, Yuan L, Zeng D, Wu K. Alteration of fractional anisotropy and mean diffusivity in glaucoma: novel results of a meta-analysis of diffusion tensor imaging studies. PLoS One. 2014;9(5):e97445. Epub 2014/05/16. doi: 10.1371/journal.pone.0097445. PubMed PMID: 24828063; PubMed Central PMCID: PMCPMC4020845.

14. Zhang YQ, Li J, Xu L, Zhang L, Wang ZC, Yang H, et al. Anterior visual pathway assessment by magnetic resonance imaging in normal-pressure glaucoma. Acta Ophthalmol. 2012;90(4):e295–302. Epub 2012/04/12. doi: 10.1111/j.1755-3768.2011.02346.x. PubMed PMID: 22489916.

15. Garaci FG, Bolacchi F, Cerulli A, Melis M, Spano A, Cedrone C, et al. Optic nerve and optic radiation neurodegeneration in patients with glaucoma: in vivo analysis with 3-T diffusion-tensor MR imaging. Radiology. 2009;252(2):496–501. Epub 2009/05/14. doi: 10.1148/radiol.2522081240. PubMed PMID: 19435941.

16. Chang ST, Xu J, Trinkaus K, Pekmezci M, Arthur SN, Song SK, et al. Optic nerve diffusion tensor imaging parameters and their correlation with optic disc topography and disease severity in adult glaucoma patients and controls. J Glaucoma. 2014;23(8):513–20. Epub 2013/05/02. doi: 10.1097/IJG.0b013e318294861d. PubMed PMID: 23632406; PubMed Central PMCID: PMCPMC3800509.

17. Wang MY, Wu K, Xu JM, Dai J, Qin W, Liu J, et al. Quantitative 3-T diffusion tensor imaging in detecting optic nerve degeneration in patients with glaucoma: association with retinal nerve fiber layer thickness and clinical severity. Neuroradiology. 2013;55(4):493–8. Epub 2013/01/30. doi: 10.1007/s00234-013-1133-1. PubMed PMID: 23358877.

18. Sidek S, Ramli N, Rahmat K, Ramli NM, Abdulrahman F, Tan LK. Glaucoma severity affects diffusion tensor imaging (DTI) parameters of the optic nerve and optic radiation. Eur J Radiol. 2014;83(8):1437–41. Epub 2014/06/09. doi: 10.1016/j.ejrad.2014.05.014. PubMed PMID: 24908588.

19. Bolacchi F, Garaci FG, Martucci A, Meschini A, Fornari M, Marziali S, et al. Differences between proximal versus distal intraorbital optic nerve diffusion tensor magnetic resonance imaging properties in glaucoma patients. Invest Ophthalmol Vis Sci. 2012;53(7):4191–6. Epub 2012/05/10. doi: 10.1167/iovs.11-9345. PubMed PMID: 22570349.

20. Allen B, Schmitt MA, Kushner BJ, Rokers B. Retinothalamic White Matter Abnormalities in Amblyopia. Invest Ophthalmol Vis Sci. 2018;59(2):921–9. Epub 2018/02/17. doi: 10.1167/iovs.17-22930. PubMed PMID: 29450539.

21. Allen B, Spiegel DP, Thompson B, Pestilli F, Rokers B. Altered white matter in early visual pathways of humans with amblyopia. Vision Research. 2015;114:48–55. doi: http://dx.doi.org/10.1016/j.visres.2014.12.021.

22. Andersson JL, Skare S, Ashburner J. How to correct susceptibility distortions in spin-echo echo-planar images: application to diffusion tensor imaging. Neuroimage. 2003;20(2):870–88. Epub 2003/10/22. doi: 10.1016/s1053-8119(03)00336-7. PubMed PMID: 14568458.

23. Smith SM, Jenkinson M, Woolrich MW, Beckmann CF, Behrens TE, Johansen-Berg H, et al. Advances in functional and structural MR image analysis and implementation as FSL. Neuroimage. 2004;23 Suppl 1:S208–19. Epub 2004/10/27. doi: 10.1016/j.neuroimage.2004.07.051. PubMed PMID: 15501092.

24. Ophthalmology AAo. ICD-10 Glaucoma Reference Guide. American Academy of Ophthalmology; 2015.

25. Basser PJ, Mattiello J, Lebihan D. Estimation of the Effective Self-Diffusion Tensor from the NMR Spin Echo. Journal of Magnetic Resonance, Series B. 1994;103(3):247–54. doi: https://doi.org/10.1006/jmrb.1994.1037.

26. Basser PJ, Jones DK. Diffusion-tensor MRI: theory, experimental design and data analysis - a technical review. NMR Biomed. 2002;15(7–8):456–67. Epub 2002/12/19. doi: 10.1002/nbm.783. PubMed PMID: 12489095.

27. Behrens TE, Woolrich MW, Jenkinson M, Johansen-Berg H, Nunes RG, Clare S, et al. Characterization and propagation of uncertainty in diffusion-weighted MR imaging. Magn Reson Med. 2003;50(5):1077–88. Epub 2003/10/31. doi: 10.1002/mrm.10609. PubMed PMID: 14587019.

28. Calamante F, Tournier JD, Jackson GD, Connelly A. Track-density imaging (TDI): super-resolution white matter imaging using whole-brain track-density mapping. Neuroimage. 2010;53(4):1233–43. Epub 2010/07/21. doi: 10.1016/j.neuroimage.2010.07.024. PubMed PMID: 20643215.

29. Conturo TE, Lori NF, Cull TS, Akbudak E, Snyder AZ, Shimony JS, et al. Tracking neuronal fiber pathways in the living human brain. Proc Natl Acad Sci U S A. 1999;96(18):10422–7. Epub 1999/09/01. PubMed PMID: 10468624; PubMed Central PMCID: PMCPMC17904.

30. Mori S, van Zijl PC. Fiber tracking: principles and strategies - a technical review. NMR Biomed. 2002;15(7–8):468–80. Epub 2002/12/19. doi: 10.1002/nbm.781. PubMed PMID: 12489096.

31. Parker GJ, Haroon HA, Wheeler-Kingshott CA. A framework for a streamline-based probabilistic index of connectivity (PICo) using a structural interpretation of MRI diffusion measurements. J Magn Reson Imaging. 2003;18(2):242–54. Epub 2003/07/29. doi: 10.1002/jmri.10350. PubMed PMID: 12884338.

32. Tournier JD, Calamante F, Connelly A. Robust determination of the fibre orientation distribution in diffusion MRI: non-negativity constrained super-resolved spherical deconvolution. Neuroimage. 2007;35(4):1459–72. Epub 2007/03/24. doi: 10.1016/j.neuroimage.2007.02.016. PubMed PMID: 17379540.

33. Tournier JD, Calamante F, Gadian DG, Connelly A. Direct estimation of the fiber orientation density function from diffusion-weighted MRI data using spherical deconvolution. Neuroimage. 2004;23(3):1176–85. Epub 2004/11/06. doi: 10.1016/j.neuroimage.2004.07.037. PubMed PMID: 15528117.

34. Tournier JD, Calamante F, Connelly A. MRtrix: Diffusion tractography in crossing fiber regions. 2012;22(1):53–66.

35. Yeatman JD, Dougherty RF, Myall NJ, Wandell BA, Feldman HM. Tract Profiles of White Matter Properties: Automating Fiber-Tract Quantification. PLOS ONE. 2012;7(11):e49790.

36. Alexander AL, Lee JE, Lazar M, Field AS. Diffusion tensor imaging of the brain. Neurotherapeutics. 2007;4(3):316–29. Epub 2007/06/30. doi: 10.1016/j.nurt.2007.05.011. PubMed PMID: 17599699; PubMed Central PMCID: PMCPMC2041910.

37. Song SK, Sun SW, Ramsbottom MJ, Chang C, Russell J, Cross AH. Dysmyelination revealed through MRI as increased radial (but unchanged axial) diffusion of water. Neuroimage. 2002;17(3):1429–36. Epub 2002/11/05. PubMed PMID: 12414282.

38. Malania M, Konrad J, Jagle H, Werner JS, Greenlee MW. Compromised Integrity of Central Visual Pathways in Patients With Macular Degeneration. Invest Ophthalmol Vis Sci. 2017;58(7):2939–47. Epub 2017/06/09. doi: 10.1167/iovs.16-21191. PubMed PMID: 28594428.

